# Brain network analysis in Parkinson’s disease patients based on graph theory

**DOI:** 10.1101/2023.02.21.529361

**Authors:** Shirin Akbari, Mohammad Reza Deevband, Amin Asgharzadeh Alvar, Emadodin Fatemi Zadeh, Hashem Rafie Tabar, Patrick Kelley, Meysam Tavakoli

## Abstract

Development of Parkinson’s disease causes functional impairment in the brain network of Parkinson’s patients. The aim of this study is to analyze brain networks of people with Parkinson’s disease based on higher resolution parcellations and newer graphical features. The topological features of brain networks were investigated in Parkinson’s patients (19 individuals) compared to healthy individuals (17 individuals) using graph theory. In addition, four different methods were used in graph formation to detect linear and nonlinear relationships between functional magnetic resonance imaging (fMRI) signals. The functional connectivity between the left precuneus and the left amygdala, as well as between the vermis 1-2 and the left temporal lobe was evaluated for the healthy and the patient groups. The difference between the healthy and patient groups was evaluated by non-parametric t-test and U-test. Based on the results, Parkinson’s patients showed a significant decrease in centrality criterion compared to healthy subjects. Furtheremore, changes in regional features of brain network were observed. There was also a significant difference between the two groups of healthy subjects and Parkinson’s patients in different areas by applying centrality criterion and the correlation coefficients. The results obtained for topological features indicate changes in the functional brain network of Parkinson’s patients. Finally, similar areas obtained by all three methods of graph formation in the evaluation of connectivity between paired regions in the brain network of Parkinson’s patients increased the reliability of the results.

## Introduction

Parkinson’s disease is the second most progressive functional neurological disorder after Alzheimer’s disease which affects nearly 1–2% people more than 60 years old worldwide [1]. The disease is initially characterized by motor symptoms, including slowed movement (bradykinesia), rigidity, body tremor, and physical condition instability (impaired posture and balance), as well as non-motor symptoms such as depression, cognitive impairments, anxiety, or sudden behavior changes [2,3]. The pathology of Parkinson’s disease is not fully understood, but there are disorders in the structure and function of the cortical and subcortical brain networks. Dopamine is a neurotransmitter which is found primarily in the neurons of the substantianigra and the frontal cortex, which are responsible for controlling motility and approach behavior. Dopamine levels for neurotransmission are found to be low in the striatum in Parkinson’s patients [4,5].

New methods in functional brain imaging provide important insights into a better understanding of the pathology of Parkinson’s disease, its treatment implications and its specific complications. These imaging studies have demonstrated physiologic dysfunction of extensive cortical areas. However, the approach in which cortical or subcortical pathology disorders interaction within and between cortical areas hasn’t done comprehensively and remains to be established [6,7]. Here, two types of functional magnetic resonance imaging (fMRI), resting-state (rs-fMRI) and in-task mode (task-fMRI), are used as non-invasive methods for psychological evaluation and are also used to discover mechanisms at the brain level, as well as to diagnose and analyze neurological diseases [8]. Over the past decade, researchers have evaluated the blood-oxygen-level-dependent (BOLD) signal in fMRI called ‘‘resting state” (rs-fMRI) studies [9,10]. This method is based on the fact that more blood is pumped into areas where the activity of the nerve cells is higher, increasing the level of oxygenated hemoglobin in the areas, which causes the BOLD signal to be produced. In effect, BOLD signals demonstrate neuronal activity in the location of the brain where the signals originate. In rs-fMRI mode, individuals are placed into the scanner in a conscious state without doing anything special. The rs-fMRI is used to examine different networks in order to show the continuous internal activity of the human brain that is affected by specific diseases. This mode is not dependent on the optimal set of conditions such as task-fMRI [11]. Therefore, researchers in clinical practice are more likely to increase the statistical power of their study by employing more people, thereby enabling them to study the functional differences in Parkinson’s disease without affecting error in the detection of specific tasks. This is important because in task-fMRI studies the activity or inactivity in motor or non-motor tasks due to abnormal brain function is not recognizable [12,13].

In recent years, graph theory has been used as a powerful tool to investigate abnormal brain network function in brain diseases such as Parkinson. Graph analysis considers the brain as a complex network of nodes linked by edges to discover the topological structure of brain networks. This method assesses the general and local characteristics of brain regions in healthy and abnormal individuals [14–18]. The subjects that include a specific phenomenon or task in Parkinson patients, such as motor, behavioral, cognitive, have been investigated in multiple studies, but these works are generally not all analyses using graph theory [12,19–21]. Furthermore, some network analysis of Parkinson patients using graph theory has only been examined in a few studies, using fMRI, structural MRI, and magnetoencephalography [17]. The combination of graph theory analysis and rs-fMRI allows extracting valuable information from the complex human brain network [22]. It has been proven using this method is associated with the development of Alzheimer’s disease [23], schizophrenia [24], depression [25]. By now, limited studies have been conducted in Parkinson’s disease using graph theoretical method or medication or disease conditions [26–28]. Although some efforts and studies have been made, less conclusive inadequate, and inconsistent results have been presented. Therefore, more investigation needs in Parkinson patients by the use of graph theory approach.

In the same direction, there are different toolboxes that were developed to study brain connectivity, including the Brain Connectivity Toolbox [29], eConnectome [30], GAT [31], CONN [32], BrainNet Viewer [33], GraphVar [34] and GRETNA [35]. Furthermore, by emerging the time-varying brain networks to identify mental illnesses [36], multiple toolboxes have also been developed to calculate dynamic functional connectivity measures [37]. While all of these works had important contributions by proving new options to build, characterize and visualize brain network topology, they need different level of computer programming experience, in such a way that their adaptation is difficult to achieve. Therefore, a reliable, streamlined, user-friendly, fast, and scalable software that deals with all aspects of network organization is still missing.

In this study, the aim is to investigate the brain connectivity features obtained from graph analysis, which was modified in the rs-fMRI brain network of Parkinson’s patients. Additionally, the brain networks of people with Parkinson’s disease is analyzed better and more accurately with variable analytic approaches, using higher-resolution parcellations as well as newer graphical features. The use of newer features may also better reflect the possible differences between the brain network of Parkinson’s patients and normal subjects.

## Materials and methods

### Design of study groups and brain network

In this work, 19 subjects with Parkinson’s disease and 17 healthy control subject were evaluated. All of the subjects were matched in terms of sex and age. The duration of study was also equal for all of them. They were accepted from data recorded in Parkinson’s progressive markers initiative (PPMI). This data were recorded in 2015 and later employed in this research in 2016. The subjects with Parkinson were observed for a 2-year period, without being completely treated. Inclusion criteria for the Parkinson patients are listed below for which the patients must hold at least two of the items: (1) Vibration in relaxed position, (2) Retardation in movements, (3) Difficulty in movements, (4) Any asymmetric vibration in relaxed position or retardation in asymmetric movements, (5) Disuse of any frequent medicine, (6) Lack of dopamine inefficiency in DaTSCAN imaging.

Inclusion criteria for healthy subjects are stated as below: (i) without history of neurological disorders, (ii) without history of Parkinson’s disease in their close family, and (iii) normal cognitively based on Montreal Cognitive Assessment. After pre-processing of fMRI signal of each subject, the whole brain was divided into 116 target regions (58 regions per brain hemisphere) using high resolution atlas. Each region has a time series which was calculated by averaging the signals of all the voxels relating to that region. The time series calculated for each region was a vector of 1 × 210 dimensions. Therefore, for each individual in the database, 116 signals with dimensions of 1 × 210 were calculated using pre-processed fMRI data, and each individual had a 210 × 116 matrix, each column representing the time series of each region. Thereafter, Pearson, Spearman, Kendall, and Copula coefficients (different types of parametric and non-parametric correlations) were calculated for each pair of time series from the selected areas and four correlation matrices were obtained for each individual with 116 × 116 dimensions.

### Recording data and preprocessing

All structural and functional images of whole brain of the individuals were taken by a 3 Tesla Siemens MR System. Structural images were recorded in a three dimensional high resolution parabolic plate by *T*_1_-weighted magnetization prepared rapid gradient-echo (MP-RAGE). fMRI data while implementing resting mode with echo-planar imaging (EPI) gradient were obtained in a session of 8 minutes and 29 seconds. These functional images contained 212 EPI volumes, each with 40 incisions in ascending order. During imaging, the sample people were resting and keeping their eyes open to stay alert. Table 1 lists the parameters for obtaining the structural and functional images used in this study.

**Table 1.** Parameters of structural and functional images.

|  | Repeating time | Eco time | Flip angle | Visibility | Voxel size | Slice thickness |
| --- | --- | --- | --- | --- | --- | --- |
| Structural images | 2300 ms | 2.98 ms | 9 degree | 256 mm | $1 \times 1 \times 1 \text{ mm}^3$ | 1 mm |
| Functional images | 2400 ms | 25 ms | 80 degree | 222 mm | $3.3 \times 3.3 \times 3.3 \text{ mm}^3$ | 3.29 mm |

Written and verbal consent was obtained from each patient selected for imaging. The evaluation was performed based on the ethical guidelines of Shahid Beheshti University of medical sciences.

For preprocessing of rs-fMRI data we performed a Matlab-based cross-platform software, CONN: Functional Connectivity Toolbox [32]. Here, the preprocessing steps included: (1) acquisition time differences correction in between slices; (2) unsupervised learning registration for motion correction during data acquisition and co-registration of the MR image to the mean EPI scans [38]; (3) Spatial normalization of the EPI images using the normalization parameters estimated from the structural image to a standard template (Montreal Neurological Institute) by DARTEL ‘normalization’; (4) image smoothing with a 4 mm full width half maximum Gaussian kernel; and (5) noise reduction by applying temporal bandpass filtering to each voxel time series. Steps 3 and 5 were added as extera parts in the CONN software.

### Graph theory and determination of regions of interest

Graph is a mathematical structure which models the connection between two or more things. It consists of nodes and edges, which are links between the nodes. The number of nodes, existence and strength of connection between them create network building blocks. Graph theory employs graphs to exemplify these individual pieces of the networks and quantify their connections to each other.

For determining regions of interest (ROI), first step in analyzing the network, is to determine the nodes of the graph which represents brain areas as a coherent patterns of anatomical or functional connections. In this research, anatomical classification of normalized volumes of subjects was employed using high resolution *T*_1_ images. For this purpose, the atlas divided the brain to 116 ROIs. First, brain grooves of each subject were determined and according to the results, three dimensional anatomical volumes (58 volumes in each brain hemisphere) were specified. The obtained anatomical areas, were based on cortical and subcortical structures and were districted with labels. The obtained network was based on the areas, therefore it indicates the microscopic structures of the brain. 116 derived areas were divided into 6 brain networks: default mode (yellow), Sensorimotor (navy), Frontal Parietal Lobe (light blue), Occipital lobe (blue), Cingulo-opercular/limbic (orange), Cerebellum (red); the brain networks are displayed by their color in Fig. 1.

**Fig 1.**
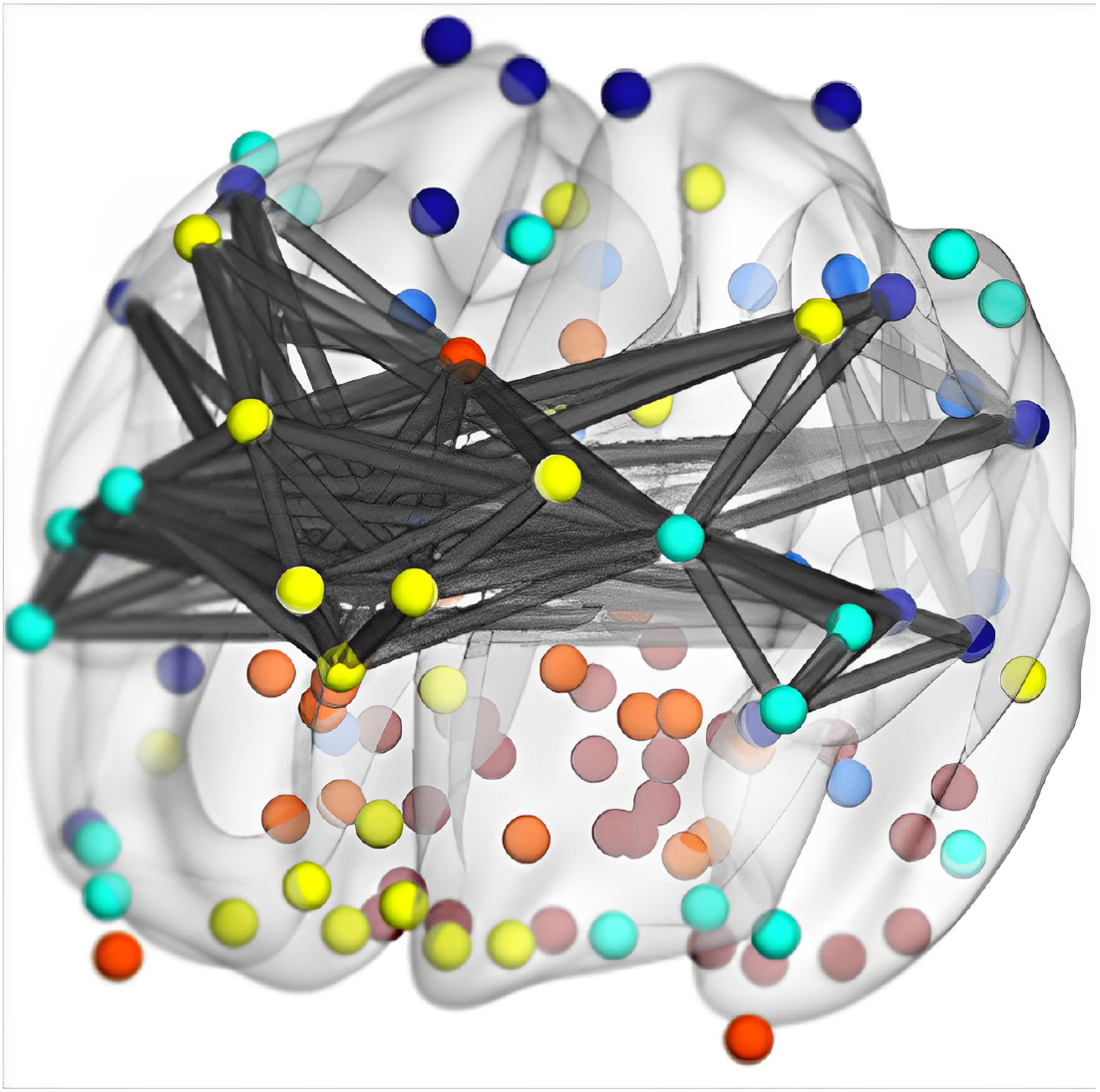
Graph nodes and surfaces. A three dimensional view of the obtained graph nodes and surfaces for Parkinson’s disease.

### Extraction of time series from target areas and correlation map

Each person had 210 volumes and each contains 40 slices. Each region contained a number of voxels and the BOLD signals of each voxel was obtained. The BOLD signal of the time series of each region was calculated from the average time series of voxels in that region. The time series obtained for each region was a vector of 1 × 210 dimensions. At this point, the target was the time series of the 116 areas. Therefore, for each individual 116 database of 210 × 1 signals is computed using pre-processed fMRI data. These time series represent the range of the activity obtained from the BOLD signal in terms of time units, and in fact describe the functional dynamics of each resting state over time for a particular region.

Slow pulsation of blood in certain brain areas describes subjects’ brain in a state in which they are doing nothing particular. These pulsations occur simultaneously in the related brain functional areas, and these areas are those even areas which are located distant from each other. These simultaneous pulsations represents neural connection between them. Functional connection is a term used for interpreting this correlation between related functional brain areas. Correlated pulsations in fMRI signal display simultaneous changes in the related functional brain areas from their BOLD signals. Evaluation of functional connection in magnetic resonance images demonstrates the intensity of brain signal alterations map. Areas having simultaneous changes in blood circulation were identified in fMRI and the map of this related functional brain areas was derived. These synchronized brain signal intensity pulsations, determines the power of neural connections between brain areas while they are doing nothing specific. Following the proposed method and all of the employed steps for analyzing the brain network, the atlas data was applied for determination of intended brain network areas and then time series of each area was derived. Lastly, formed correlation map of brain network was elucidated by applying correlation coefficients.

## Results

### Network centralization

The centralization value characterizes the influence of the nodes; in other words, it is used to determine important areas of the brain, represented by the connections of the network nodes, that function differently between healthy brains and those afflicted with Parkinson’s disease. Network centralization via centrality indices is calculated from each correlation coefficient, i.e. the edges, upon which the significance of each node can be numerically ascertained. Since the number of individuals was 17 in healthy group and 19 in patient group, by calculating the general network centrality for each person, a 19 × 1 vector for the patients set and a 17 × 1 vector for the healthy individuals set were applied to each centrality index. By calculating mean centralization in the healthy and patient groups and by applying each centrality index, it was observed that the brain network centralization of the healthy group yielded larger values than the patient group. Fig. 2 shows the results of brain network centralization obtained from different correlation coefficients in healthy individuals and patients using degree centrality, eigenvectors, Katz and descending methods. To obtain distribution of each vector, JBTEST test was applied. For those areas with normal distribution, t-test and for those without normal distribution, U-test was applied. Comparison of the values obtained from centralization calculation revealed no significant difference between the healthy and the patient groups.

**Fig 2.**
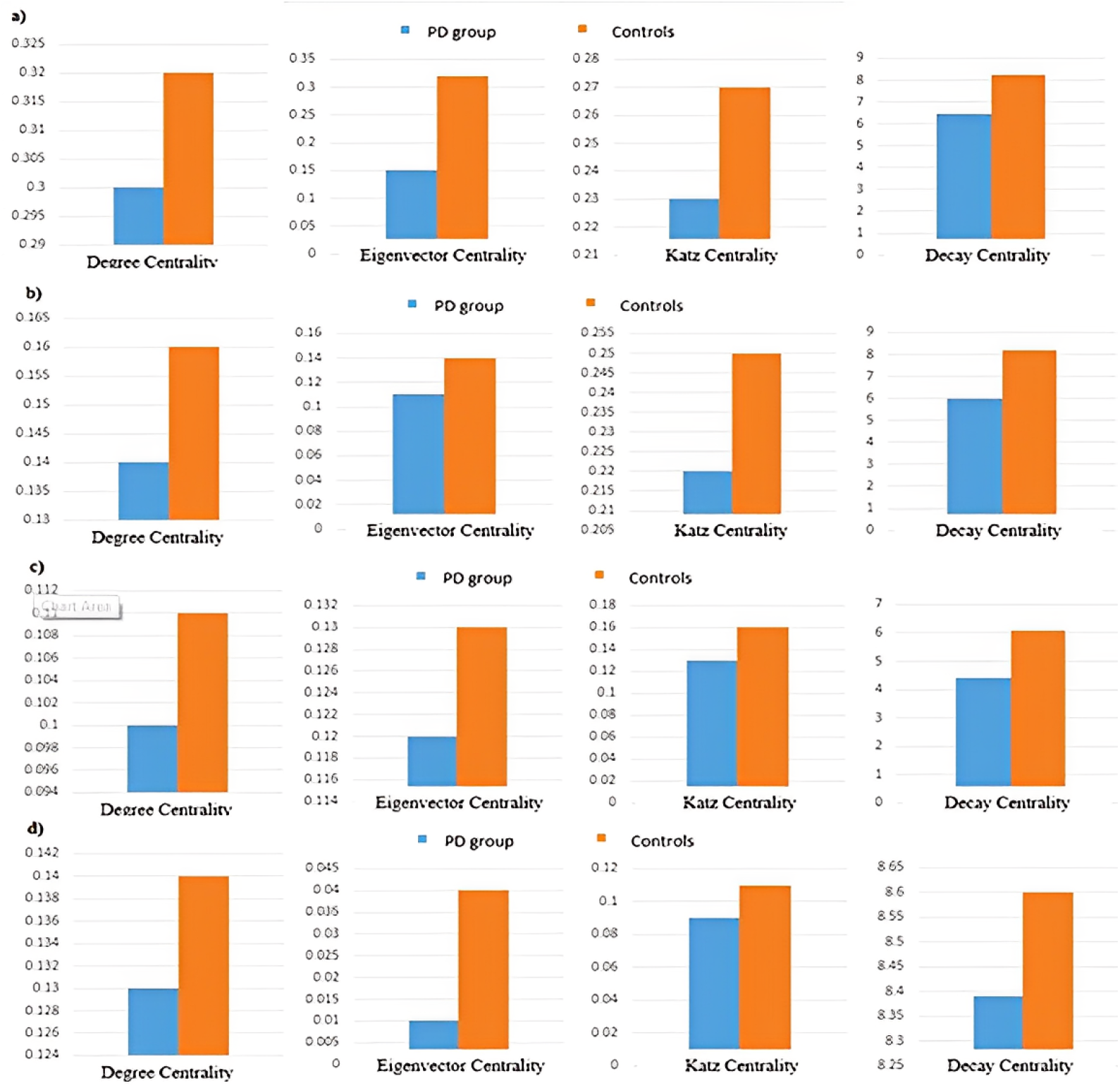
Results of brain network centralization. Centrality values calculated by degree centrality, eigenvectors, Katz and descending measures in brain network obtained by Pearson correlation coefficient (a), Spearman correlation coefficient (b), Kendall correlation coefficient (c), and Copula correlation coefficient (d).

### Brain network local criteria

In this section, three features of centrality containing degree centrality, eigenvector, and closeness were used. Here, the degree centrality comes up with information only in the local popularity. It determines which nodes can quickly and directly propagated in an area. In degree centrality the significance of a node is considered with the number of its neighbors [39]. The eigenvector centrality provides global information that a predominant node is connected to other well-connected nodes in a network [40]. At last, closeness measurment specifies the global “close” distance of a node to other nodes [39].

Three characteristics from studies of brain network changes in Parkinson’s patients by using Katz, and Descending centrality were first applied to fMRI data from these patients to find areas with significant differences between healthy and patient groups. Here, Katz centrality uses an eigenvector by weighting the connections involving more highly connected nodes higher than those from lesser connected nodes when considering the makeup of the shortest paths which pass through a given node [41,42]. To investigate the effect of the characteristics on making a significant difference between healthy and patient groups, U-test (for data without normal distribution) and t-test (for data with normal distribution) were used. The mean time series of the signals of all voxels for each region was a vector of 210 × 1 dimensions. Since the number of divided zones was 116, a 210 × 116 matrix was obtained for each person, each column representing the time series for each zone. For each individual in both healthy and patient groups, the features were applied on the obtained correlation matrices. Since the atlas divides brain into 116 zones, the correlation matrix was 116 × 116. The relationship between brain regions was determined by correlation analysis. For this purpose, four correlation methods, Pearson, Spearman, Kendall, and Copula correlation coefficient, were used to investigate the relationship between brain regions and the effects of each feature on two groups of patients and healthy groups.

### Degree centrality, eigenvector and closeness

Using Pearson correlation coefficient for applying degree centrality, 21 areas with significant differences between patient and healthy groups were obtained. Furthermore, the results of eigenvector centrality revealed 9 areas and closeness centrality revealed 11 areas with significant differences between the two groups of patients and healthy ones. Common areas identified in the eigenvector centrality and degree centrality measures include right rectus, right posterior cingulum and right inferior temporal. Two measures of degree centrality and closeness shared the right amygdala. Table 2 shows significant changes in different brain regions in examining the degree centrality, eigenvector and closeness between controls and Parkinson’s patients. Table 3 presents an example of previous studies using the degree centrality and Betweenness centrality measures. As it can be seen, in this study degree centrality identified 26 areas and Betweenness centrality identified 12 areas with a significant difference between the two groups. Based on the results, some of the obtained results are similar to those obtained in this study with applying degree centrality measure. These areas include amygdala and putamen, which are areas of the basal ganglia. According to the study, the centrality index of some areas of the brain, such as bilateral pallidum, bilateral amygdala, right medial superior frontal gyrus, and certain areas of the parietal lobes, were higher in Parkinson’s patients. Furthermore, some areas in the brain of Parkinson’s patients, such as left motor supplemental region, caudate nucleus, precentral gyrus, right frontal gyrus, fusiform gyrus and occipital gyrus have shown decreased nodal centrality compared to the healthy individuals.

**Table 2.** Areas obtained in changes between healthy and desease groups based on centrality criteria (p < 0.05).

| Anatomical Region | Eigenvector centrality<br>p-values | Degree Centrality<br>p-values | Closeness centrality<br>p-values |
| --- | --- | --- | --- |
| Frontal Inf Oper L | > 0.05 | 0.009 | > 0.05 |
| Frontal Inf Tri L | > 0.05 | 0.043 | > 0.05 |
| Frontal Inf Orb L | > 0.05 | 0.031 | > 0.05 |
| Supp Motor Area L | > 0.05 | 0.011 | > 0.05 |
| Rectus R | > 0.05 | 0.043 | > 0.05 |
| Cingulum Post L | > 0.05 | 0.027 | > 0.05 |
| Cingulum Mid L | > 0.05 | > 0.05 | 0.049 |
| Cingulum Post R | > 0.05 | > 0.05 | 0.042 |
| Hippocampus L | 0.046 | > 0.05 | > 0.05 |
| Hippocampus R | 0.027 | > 0.05 | > 0.05 |
| Amygdala R | > 0.05 | 0.005 | 0.017 |
| Lingual R | 0.036 | > 0.05 | > 0.05 |
| Occipital Mid L | > 0.05 | 0.034 | > 0.05 |
| Occipital Mid R | > 0.05 | > 0.05 | 0.022 |
| Fusiform L | > 0.05 | 0.049 | > 0.05 |
| Paracentral Lobule L | > 0.05 | 0.036 | > 0.05 |
| Paracentral Lobule R | > 0.05 | > 0.05 | 0.002 |
| Putamen L | > 0.05 | 0.005 | > 0.05 |
| Putamen R | > 0.05 | 0.031 | > 0.05 |
| Pallidum L | > 0.05 | > 0.05 | 0.017 |
| Heschl L | > 0.05 | 0.029 | > 0.05 |
| Temporal Pole Sup R | > 0.05 | > 0.05 | 0.011 |
| Temporal Mid L | > 0.05 | > 0.05 | 0.007 |
| Cerebelum Crus L | > 0.05 | > 0.05 | 0.036 |
| Cerebelum 8 L | 0.043 | > 0.05 | > 0.05 |
| Cerebelum 8 R | 0.029 | > 0.05 | > 0.05 |
| Cerebelum 9 R | 0.031 | > 0.05 | > 0.05 |
| Temporal Inf R | 0.023 | 0.023 | > 0.05 |
| Cerebelum 3 R | > 0.05 | 0.021 | > 0.05 |
| Cerebelum 4 5 L | > 0.05 | 0.011 | > 0.05 |
| Cerebelum 4 5 R | > 0.05 | 0.003 | > 0.05 |
| Cerebelum 6 L | > 0.05 | 0.049 | > 0.05 |
| Cerebelum 6 R | > 0.05 | 0.046 | > 0.05 |
| Vermis 1 2 | > 0.05 | 0.046 | > 0.05 |
| Vermis 3 | > 0.05 | > 0.05 | 0.033 |
| Vermis 4 5 | > 0.05 | 0.046 | > 0.05 |
| Vermis 6 | > 0.05 | 0.046 | > 0.05 |
| Vermis 10 | > 0.05 | > 0.05 | 0.036 |

**Table 3.**
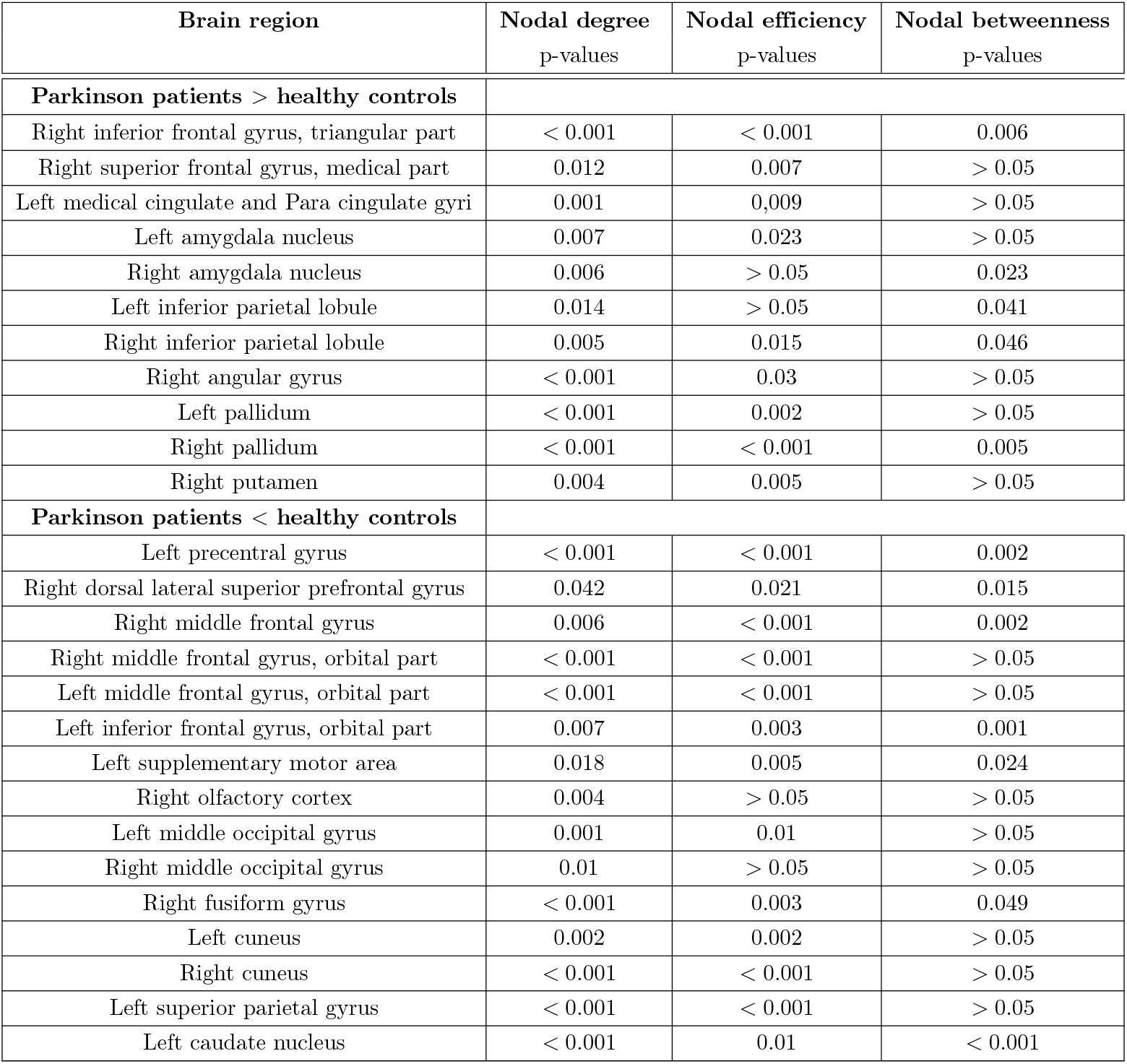
Areas with abnormal results in Parkinson’s disease in the previous studies.

### Katz centrality

With the use of Pearson correlation coeffients, we implementing the Katz centrality feature and obtained 25 zones with significant difference between patient and healthy groups. By using Spearman correlation coefficient instead for this feature, 39 areas were obtained with significant difference between the two groups. The results of Kendall correlation coefficient as parameters in the Katz centrality feature showed 41 areas with significant difference between the patient and healthy groups. The ratios were obtained by applying Copula correlation coefficient in this feature to 18 regions with a significant difference between the two groups. By examining the results of the features with the use of Pearson, Spearman and Kendall correlation coefficients, 23 common areas were obtained. By using the Copula correlation coefficient, 9 common areas with other correlation coefficients were obtained. Using the Pearson, Spearman and Kendall correlation coefficients, the putamen and amygdala regions, which are parts of the basal ganglia, were detected with very low p-values, indicating a lower error coefficient in detection of significant differences between the two groups. Using the Copula correlation coefficient, Significant differences between the areas of the left upper anterior apical and the left upper anterior triangular with p-values of 0.0004 and 0.005 were also observed, respectively.

## Descending centrality

According to the observations, 13 zones were obtained by using the Pearson correlation coefficient with applying Decay centrality feature, showing significant difference between patient and healthy groups. By using Spearman correlation coefficient with applying Decay centrality feature, 35 areas were obtained with significant difference between the patient and healthy groups. The results of Kendall correlation coefficient with applying Decay centrality feature showed 33 areas with significant difference between patient and healthy groups. By using the Copula correlation coefficient with applying Decay centrality feature, 18 zones were obtained with significant difference between the patient and healthy groups. In Tables 4, 5, 6 and 7 the results are listed for each of the features that showed a significant difference between the regions with the Pearson, Spearman, Kendall, and Copula correlation coefficients (*p* < 0.05). The specified areas in these tables, show the common results of applying these correlation coefficients by using each feature. Accordingly, 23 zones with applying the Katz Centrality feature and 4 common zones with applying the Decay Centrality feature were obtained. The small P-values in these table, indicate low coefficient uncertainties in detection of significant differences. By applying the Coppola correlation coefficient, with applying the Katz Centrality feature, 9 zones, and with the Decay Centrality feature, 2 common zones were obtained with other correlation coefficients. Using the three Pearson, Spearman, and Kendall correlation coefficients, some important areas in Parkinson’s disease such as left and right cuneus, left and right putamen and left caudate were detected with very low p-values, indicating low coefficient uncertainties in the diagnosis of significant difference between the two groups.

**Table 4.**
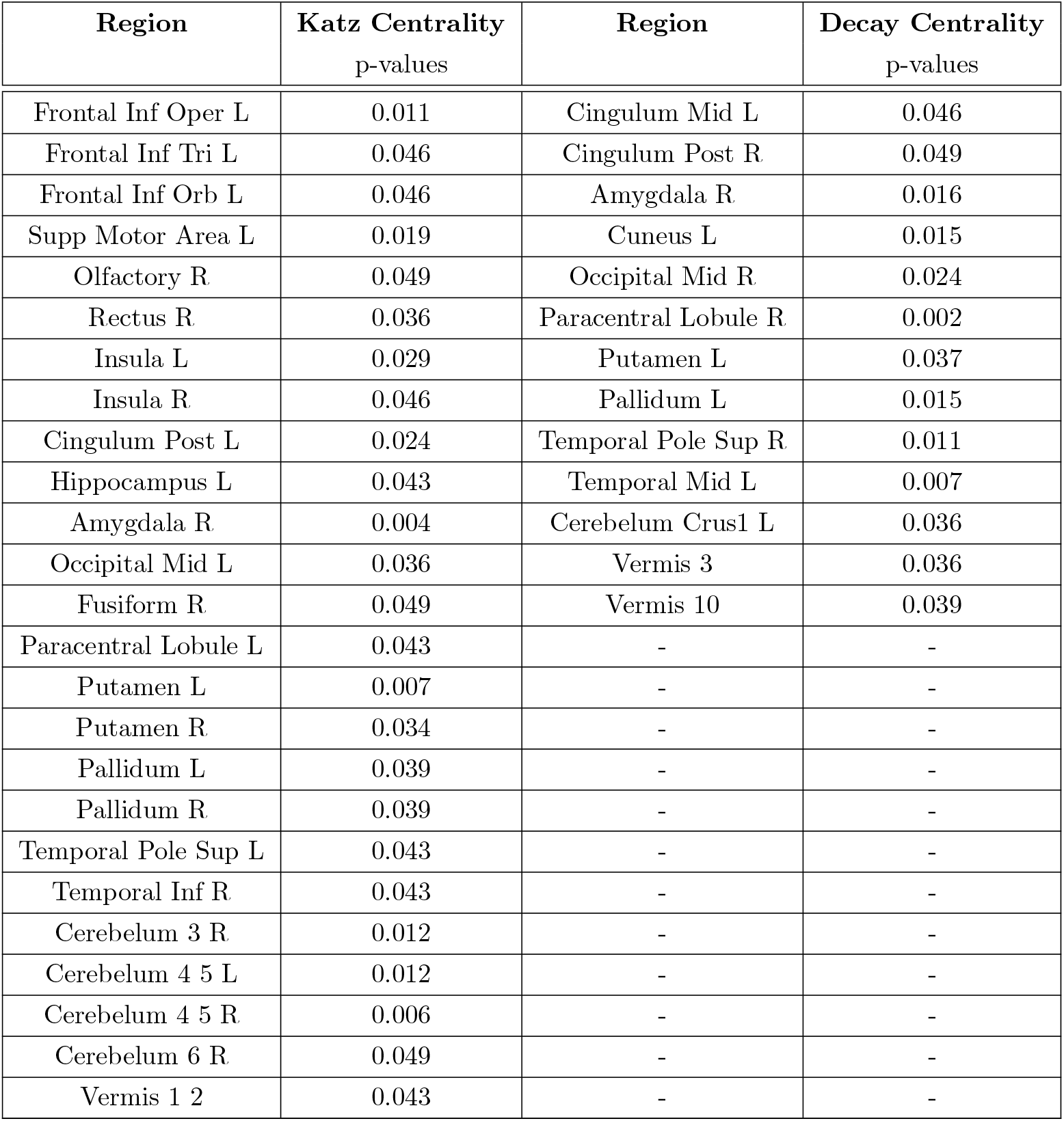
Areas with significant outcomes in Parkinson’s patients compared with healthy individuals using Pearson correlation coefficient.

**Table 5.** Areas with significant outcomes in Parkinson’s patients compared with healthy individuals using Spearman correlation coefficient.

| Region | Katz Centrality<br>p-values | Region | Decay Centrality<br>p-values |
| --- | --- | --- | --- |
| Frontal Inf Oper L | 0.008 | Frontal Mid L | 0.039 |
| Frontal Inf Tri L | 0.027 | Frontal Inf Oper R | 0.019 |
| Frontal Inf Orb L | 0.024 | Supp Motor Area L | 0.043 |
| Frontal Inf Orb R | 0.039 | Supp Motor Area R | 0.034 |
| Rolandic Oper L | 0.046 | Frontal Sup Medial L | 0.043 |
| Supp Motor Area L | 0.011 | Frontal Med Orb R | 0.024 |
| Olfactory R | 0.049 | Rectus R | 0.034 |
| Rectus R | 0.049 | Cingulum Mid L | 0.039 |
| Insula L | 0.024 | ParaHippocampal R | 0.029 |
| Insula R | 0.031 | Amygdala L | 0.039 |
| Cingulum Mid L | 0.049 | Calcarine L | 0.027 |
| Cingulum Post L | 0.027 | Cuneus L | 0.004 |
| Hippocampus L | 0.034 | Cuneus R | 0.009 |
| Hippocampus R | 0.034 | Occipital Mid L | 0.049 |
| ParaHippocampal R | 0.024 | Occipital Mid R | 0.030 |
| Amygdala R | 0.001 | Fusiform L | 0.009 |
| Occipital Sup L | 0.036 | Fusiform R | 0.033 |
| Occipital Mid L | 0.046 | Postcentral R 7 | 0.049 |
| Occipital Inf L | 0.039 | SupraMarginal L | 0.046 |
| Occipital Inf R | 0.049 | Precuneus R | 0.024 |
| Fusiform L | 0.034 | Caudate L | 0.007 |
| Fusiform R | 0.046 | Putamen L | 0.007 |
| Paracentral Lobule L | 0.031 | Putamen R | 0.027 |
| Caudate L | 0.029 | Pallidum R | 0.021 |
| Putamen L | 0.007 | Heschl R | 0.027 |
| Putamen R | 0.015 | Temporal Pole Sup L | 0.046 |
| Pallidum L | 0.012 | Temporal Mid L | 0.049 |
| Pallidum R | 0.031 | Temporal Pole Mid R | 0.023 |
| Temporal Sup R | 0.039 | Temporal Inf L | 0.049 |
| Temporal Pole Sup L | 0.034 | Cerebelum 4 5 L | 0.013 |
| Temporal Pole Sup R | 0.046 | Cerebelum 7b L | 0.046 |
| Temporal Mid R | 0.043 | Vermis 1 2 | 0.002 |
| Temporal Inf R | 0.024 | Vermis 3 | 0.024 |

**Table 6.** Areas with significant outcomes in Parkinson’s patients compared with healthy individuals using Kendall correlation coefficient.

| Region | Katz Centrality<br>p-values | Region | Decay Centrality<br>p-values |
| --- | --- | --- | --- |
| Frontal Inf Oper L | 0.006 | Supp Motor Area L | 0.031 |
| Frontal Inf Tri L | 0.024 | Olfactory L | 0.023 |
| Frontal Inf Orb L | 0.021 | Frontal Med Orb R | 0.032 |
| Frontal Inf Orb R | 0.046 | Rectus R | 0.043 |
| Rolandic Oper L | 0.049 | Insula R | 0.017 |
| Supp Motor Area L | 0.013 | Cingulum Post R | 0.046 |
| Olfactory R | 0.043 | Para Hippocampal R | 0.033 |
| Insula L | 0.027 | Cuneus L | 0.005 |
| Insula R | 0.034 | Cuneus R | 0.016 |
| Cingulum Mid L | 0.049 | Occipital Mid R | 0.044 |
| Cingulum Post L | 0.023 | Occipital Inf R | 0.041 |
| Hippocampus L | 0.034 | Angular R | 0.031 |
| Hippocampus R | 0.034 | Paracentral Lobule L | 0.025 |
| Para Hippocampal R | 0.023 | Caudate L | 0.006 |
| Amygdala L | 0.049 | Caudate R | 0.034 |
| Amygdala R | 0.001 | Putamen L | 0.021 |
| Occipital Sup L | 0.039 | Putamen R | 0.043 |
| Occipital Mid L | 0.043 | Pallidum R | 0.012 |
| Occipital Inf L | 0.049 | Temporal Sup L | 0.019 |
| Occipital Inf R | 0.049 | Temporal Pole Sup L | 0.024 |
| Fusiform L | 0.039 | Temporal Pole Sup R | 0.049 |
| Fusiform R | 0.034 | Temporal Mid R | 0.043 |
| Paracentral Lobule L | 0.021 | Temporal Pole Mid R | 0.031 |
| Caudate L | 0.031 | Temporal Inf L | 0.021 |
| Putamen L | 0.005 | Temporal Inf R | 0.006 |
| Putamen R | 0.015 | Cerebelum 4 5 L | 0.022 |
| Pallidum L | 0.015 | Cerebelum 4 5 R | 0.016 |
| Pallidum R | 0.034 | Cerebelum 6 R | 0.046 |
| Heschl L | 0.019 | Cerebelum 7b R | 0.039 |
| Temporal Sup R | 0.043 | Vermis 1 2 | 0.002 |
| Temporal Pole Sup L | 0.029 | Vermis 3 | 0.012 |
| Temporal Pole Sup R | 0.043 | Vermis 6 | 0.032 |
| Temporal Mid R | 0.049 | Vermis 9 | 0.031 |

**Table 7.**
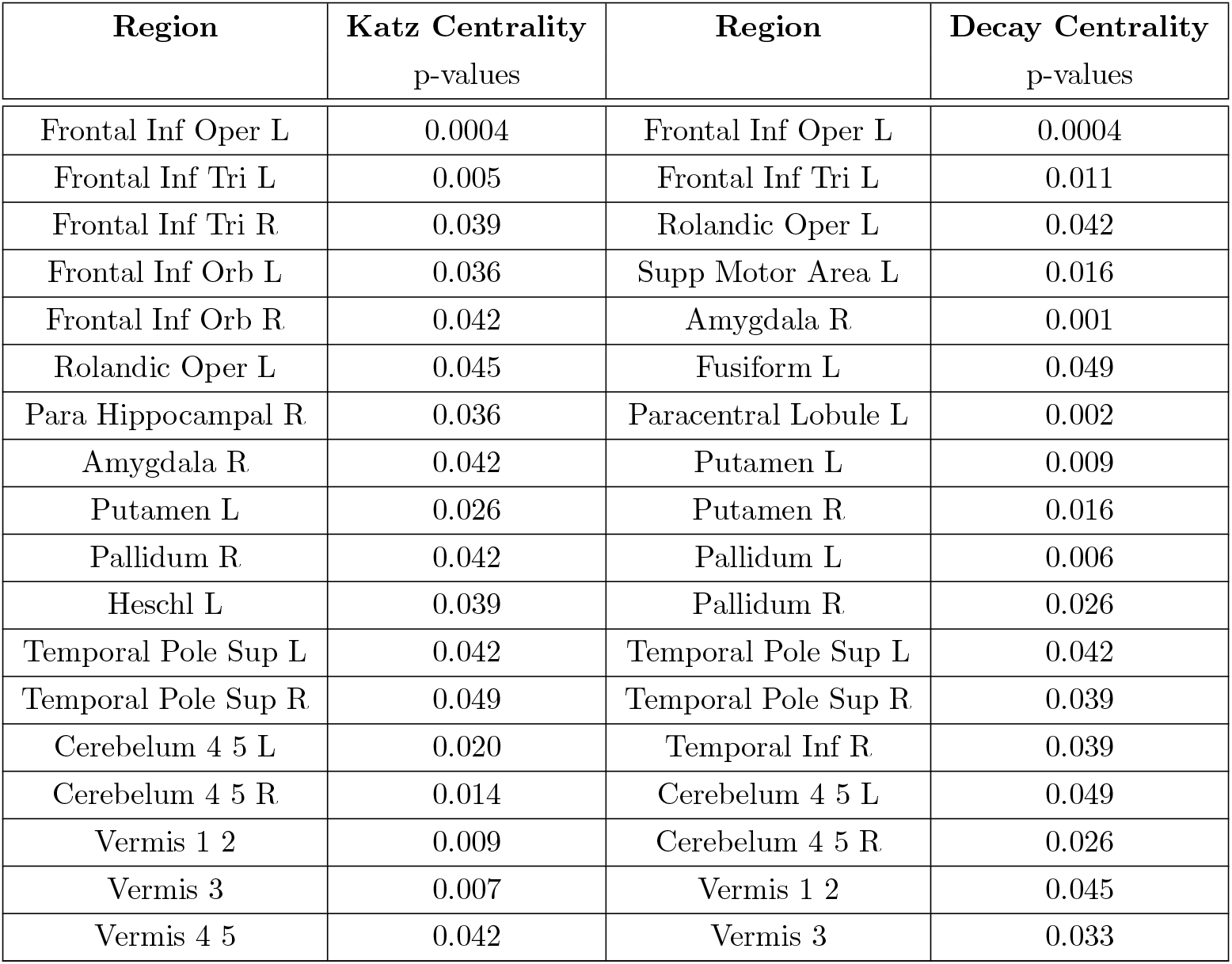
Areas with significant results in Parkinson’s patients compared with healthy individuals using Copula correlation coefficient.

| Region | Katz Centrality<br>p-values | Region | Decay Centrality<br>p-values |
| --- | --- | --- | --- |
| Frontal Inf Oper L | 0.0004 | Frontal Inf Oper L | 0.0004 |
| Frontal Inf Tri L | 0.005 | Frontal Inf Tri L | 0.011 |
| Frontal Inf Tri R | 0.039 | Rolandic Oper L | 0.042 |
| Frontal Inf Orb L | 0.036 | Supp Motor Area L | 0.016 |
| Frontal Inf Orb R | 0.042 | Amygdala R | 0.001 |
| Rolandic Oper L | 0.045 | Fusiform L | 0.049 |
| Para Hippocampal R | 0.036 | Paracentral Lobule L | 0.002 |
| Amygdala R | 0.042 | Putamen L | 0.009 |
| Putamen L | 0.026 | Putamen R | 0.016 |
| Pallidum R | 0.042 | Pallidum L | 0.006 |
| Heschl L | 0.039 | Pallidum R | 0.026 |
| Temporal Pole Sup L | 0.042 | Temporal Pole Sup L | 0.042 |
| Temporal Pole Sup R | 0.049 | Temporal Pole Sup R | 0.039 |
| Cerebelum 4 5 L | 0.020 | Temporal Inf R | 0.039 |
| Cerebelum 4 5 R | 0.014 | Cerebelum 4 5 L | 0.049 |
| Vermis 1 2 | 0.009 | Cerebelum 4 5 R | 0.026 |
| Vermis 3 | 0.007 | Vermis 1 2 | 0.045 |
| Vermis 4 5 | 0.042 | Vermis 3 | 0.033 |

### Connection between pairs of regions

The results of the evaluation of the relationship between the pairs of brain regions are reported in Table 5. According to the results, by using Pearson correlation coefficient, 665 zones with significant difference with p < 0.05 were found. For the purpose of better evaluation, the confidence coefficient was increased and this difference was considered with p < 0.001. Accordingly, 9 areas had significant differences. Using Spearman correlation coefficient, 567 zones were found to have a significant difference with p < 0.05. To evaluate better, the confidence coefficient was increased and this difference was considered with p < 0.001. Accordingly, 6 areas had significant differences. Using Kendall correlation coefficient, 570 areas with *p* < 0.05 were significantly different. For the purpose of better evaluation, the coefficient of confidence was increased and this difference was considered as p < 0.001. Accordingly, five areas had significant differences. Using the Copula correlation coefficient, 6793 areas with p < 0.05 were significantly different. To evaluate better, the confidence coefficient was increased and this difference was considered with p < 0.001. Accordingly, 521 areas were significantly different.

Investigation of similar results obtained from all three methods of graph formation increased the reliability of the results. For example, the functional connectivity between the left periconius and left amygdala regions were significantly different. There was also a significant difference between Vermis 1-2 and left temporal lobe with p < 0.001 by using three methods between the healthy and the patient groups.

The findings obtained from the analysis of brain network graphs are detailed in this study. To compare the brain network of the healthy and patient groups, t-test and U-test were used. Subsequently, the results of the overall brain network changes in patients with Parkinson’s were expressed by examining the general criteria of the graph in the name of concentration. By using the general features of the graph, different areas of the brain network revealed a significant difference between the healthy and patient groups, some of which were important areas of the basal ganglia. Finally, there were significant differences between the two groups in evaluating the association of the paired areas and the important areas were reported. Each of the general and local features and the relationships between the pair region were investigated using different graph formation methods (Pearson, Spearman, Kendall and Copula correlation coefficients).

## Discussion

In this study, the topological features of functional brain networks were examined in Parkinson’s patients and compared with a control set of healthy patients. Accordingly, changes of whole brain network were evaluated by fMRI data in Parkinson’s patients and their regional changes were evaluated by different graph indices. Gottlich et al. [27], Skidmore et al. [26] and Baggio et al. [17] applied graph theory for brain network analysis using fMRI in their studies. By different correlation coefficients, undirected weighted networks consisting of equal surfaces and nodes were formed. Based on the results, the graph theory-based method proved the modified brain network structure in Parkinson’s patients.

Some previous studies such as studies conducted by Olde Dubbelink et al. [43], Gottlich et al. [27], Baggio et al. [17], Skidmore et al. [26] and Utianski et al. [7] have investigated the general features and changes of the brain network of Parkinson’s patients, including clustering coefficient and small-world network. The current study examines brain networks of individuals by using the centralization feature for the first time. The centralization feature of brain network obtained from each correlation coefficient was evaluated in healthy and patient groups using degree centrality, eigenvector, Katz and Decay criteria. According to the observations, within all of the methods used for graph formation, including Pearson, Spearman, Kendall and Coppola correlation coefficients, the centralization feature in Parkinson’s patients was decreased. This decrease indicates that the overall brain structure of Parkinson’s patients is impaired. The decrease in centralization feature demonstrates that in the patient’s brain network (in comparison with healthy individuals), centrality criterion has decreased in the most central node has decreased compared to other nodes of the network. According to this definition, the centralized structure of the patients overall brain network was altered and defected. This means that the stiffness of the overall brain network around the central node of the network, called the focal node, has decreased and the brain network has become more dispersed.

The nodal indices used included degree centrality, eigenvector and closeness. Three local indices of graphs, including Katz centrality, and decay centrality were also used for the first time to investigate changes in the brain network of Parkinson’s patients. The different regions obtained are parts of the basal ganglia which play an important role in Parkinson’s disease. Additionally, in different areas of the brain, these indices either decreased or increased. For example, the degree centrality was increased in the right and left pallidum areas.

In Parkinson’s disease, a significant portion of the dopaminergic neurons in the substantianigra are destroyed by dopamine deficiency in the basal ganglia. This defect causes functional changes affecting all areas of the basal ganglia in Parkinson’s patients’ brain network. The basal ganglia circuit interferes with the function of the brain cortex and thalamus and modulates movements [44]. According to experimental studies, basal ganglia exert a deterrent effect on some motor systems and release of this inhibitory agent causes a motor system to function. It has been demonstrated that there are changes in the function of basal ganglia circuit in Parkinson’s patients [45, 46]. Therefore, the results of this study are approved and indicate a disorder of the basal ganglia in Parkinson’s patients.

Changes in functional connections between pairs of brain regions were investigated and many changes were observed in these connections. The changes of functional connections were calculated by the use of three different graph formation methods (Pearson, Spearman and Kendall correlation coefficients). Similar results which were obtained by all the three methods of graph formation support the reliability of the results. For example, there was a significant change in functional connection between the left precuneus and the left amygdala. There was also a significant difference between Vermis 1-2 and left temporal lobe with p < 0.001 in all the three methods between the healthy and the patient groups.

## Conclusion

Parkinson’s is a prevalent neurodegenerative disorder. Changes in the brain communication network is considered as one of the most important characteristics/effects of this disease. Our results show the application of graph theory in analyzing brain network of Parkinson’s patients. Centralization feature is decreased in Parkinson’s patients’ brain network based on the four criteria of centrality, eigenvector, Katz and descent. Additionally, their brain network is more dispersed and the density around the focal node is decreased. This impairment causes functional changes in basal ganglia of brain network of Parkinson’s patients. The results of the present study demonstrate defects in basal ganglia circuitry in Parkinson’s patients. Furthermore, the results show a significant change in functional connectivity among paired areas of the brain in the patients. More studies are needed to develop new methods of applying graph theory in order to analyze the brain network of Parkinson’s patients.

